# Does ‘information control the living state’?

**DOI:** 10.1101/003186

**Authors:** Rodrick Wallace

## Abstract

We generalize the recently-uncovered Data Rate Theorem in the context of cognitive systems having a ‘dual’ information source, including those of the living state that is particularly characterized by cognition at every scale and level of organization. The unification of information theory and control theory via the Data Rate Theorem is not additive, but synergistic, generating new statistical tools that greatly constrain the possible dynamics of that state. Thus, in addition to providing novel conceptual approaches, this emerging body of theory permits construction of models that, like those of regression analysis, can provide benchmarks against which to compare experimental or observational data.

## 1 Introduction

The keynote session of the 2014 Gordon Research Conferences section on Complex Adaptive Matter, ‘How Can Information Control Matter?’ by Goldenfeld and Ramaswamy, raises a question that has been the focus of intense debate since the end of World War II. Information theory and control theory both emerged from the technological cauldron of that conflict, engaging seemingly separate problems in communication and system regulation. The Shannon Coding and Source Coding Theorems, and the Rate Distortion Theorem, provide statistical constraints on communication, while the Bode Integral Theorem constrains system control – noise energy suppressed in one frequency range emerges in another. The two disciplines remained largely separate over the succeeding half-century.

Beginning in the late 1990’s, however, a series of studies deliniated a central relation between information theory and control theory, a direct extension of Bode’s result known now as the Data Rate Theorem that, at least partially, addresses Goldenfeld and Ramaswamy’s question. Here we will explore in more detail the implications of the unification of information and control theories for understanding the dynamics of the living state.

The underlying conceit is that living systems are cognitive at every scale and level of organization, and that many cognitive phenomena can be represented by ‘dual’ information sources constrained by the four basic theorems (e.g., Wallace, 2012a, 2014a; Wallace and Wallace, 2013). These ideas, of course, are quite old, first articulated by Maturana and Varela (1980). Wallace (2014a) and Wallace and Wallace (2013) describe gene expression, the immune system, tumor control, wound healing, animal consciousness, sociocultural cognition, and other biological processes, as examples. Wallace (2012b), in fact, provides a detailed exploration of the glycan/lectin cell surface ‘kelp bed’ as a cognitive system that necessarily rivals high order neural phenomena in its sophistication – the argument is surprisingly direct. In a sense, then, the question asked in the title to this paper is something of a red herring, reflecting a set of popular misconceptions about the relation between information and life, since information dynamics are so deeply convoluted with life itself.

We will, ultimately, expand the perspective via an ‘obvious’ generalization of the Data Rate Theorem.

### 2 The Data-Rate Theorem

The data-rate theorem, based on the Bode integral theorem for linear control systems (e.g., Yu and Mehta, 2010; Kitano, 2007; Csete and Doyle, 2002), describes the stability of linear feedback control under data rate constraints (e.g., Mitter, 2001; Tatikonda and Mitter, 2004; Sahai, 2004; Sahai and Mitter, 2006; Minero et al., 2009; Nair et al., 2007; You and Xie, 2013). Given a noise-free data link between a discrete linear plant and its controller, unstable modes can be stabilized only if the feedback data rate ***ℋ*** is greater than the rate of ‘topological information’ generated by the unstable system. For the simplest incarnation, if the linear matrix equation of the plant is of the form *x*_*t*+1_ = **A***x*_*t*_ + *…*, where *x*_*t*_ is the n-dimensional state vector at time *t*, then the necessary condition for stabilizability is

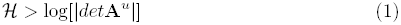

where *det* is the determinant and **A**^*u*^ is the decoupled unstable component of **A**, i.e., the part having eigenvalues ≥ 1.

There is, then, a critical positive data rate below which there does not exist any quantization and control scheme able to stabilize an unstable (linear) feedback system.

This result, and its variations, are as fundamental as the Shannon Coding and Source Coding Theorems, and the Rate Distortion Theorem (Cover and Thomas, 2006; Ash, 1990; Khinchin, 1957).

It is possible to significantly extend the argument. The essential analytic tool will be an analog to Pettini’s (2007) ‘topological hypothesis’ – a version of Landau’s spontaneous symmetry breaking insight for physical systems (Landau and Lifshitz, 2007) – which infers that punctuated events often involve a change in the topology of an underlying configuration space, and the observed singularities in the measures of interest can be interpreted as a ‘shadow’ of major topological change happening at a more basic level. The tool for the study of such topological changes is Morse Theory (Pettini, 2007; Matsumoto, 2002).

The first step is a recapitulation of an approach to cognition using the asymptotic limit theorems of information theory (Wallace 2000, 2005, 2007, 2012a, 2014a).

### 3 Cognition as an information source

Atlan and Cohen (1998) argue that the essence of cognition involves comparison of a perceived signal with an internal, learned or inherited picture of the world, and then choice of one response from a much larger repertoire of possible responses. That is, cognitive pattern recognition-and-response proceeds by an algorithmic combination of an incoming external sensory signal with an internal ongoing activity – incorporating the internalized picture of the world – and triggering an appropriate action based on a decision that the pattern of sensory activity requires a response.

Incoming sensory input is thus mixed in an unspecified but systematic manner with internal ongoing activity to create a path of combined signals *x* = (*a*_0_*, a*_1_*, …, a_n_*, …). Each *a*_*k*_ thus represents some functional composition of the internal and the external. An application of this perspective to a standard neural network is given in Wallace (2005, p.34).

This path is fed into some unspecified ‘decision function’, *h*, generating an output *h*(*x*) that is an element of one of two disjoint sets *B*_0_ and *B*_1_ of possible system responses. Let

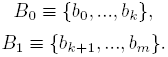

Assume a graded response, supposing that if

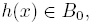

the pattern is not recognized, and if

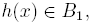

the pattern is recognized, and some action *b*_*j*_, *k* + 1 ≤ *j* ≤ *m* takes place.

Interest focuses on paths *x* triggering pattern recognition-and-response: given a fixed initial state *a*_0_, examine all possible subsequent paths *x* beginning with *a*_0_ and leading to the event *h*(*x*) ∊ *B*_1_. Thus *h*(*a*_0_*, …, a_j_*) ∊ *B*_0_ for all 0 ≤ *j < m*, but *h*(*a*_0_*, …, a*_*m*_) ∊ *B*_1_.

For each positive integer *n*, take *N*(*n*) as the number of high probability paths of length *n* that begin with some particular *a*_0_ and lead to the condition *h*(*x*) ∊ *B*_1_. Call such paths ‘meaningful’, assuming that *N*(*n*) will be considerably less than the number of all possible paths of length *n* leading from *a*_0_ to the condition *h*(*x*) ∊ *B*_1_.

Identification of the ‘alphabet’ of the states *a*_*j*_, *B*_*k*_ may depend on the proper system coarse graining in the sense of symbolic dynamics (e.g., Beck and Schlogl, 1993).

Combining algorithm, the form of the function *h*, and the details of grammar and syntax, are all unspecified in this model. The assumption permitting inference on necessary conditions constrained by the asymptotic limit theorems of information theory is that the finite limit *H* ≡ lim_*n*→∞_ log[*N*(*n*)]*/n* both exists and is independent of the path *x*. Again, *N*(*n*) is the number of high probability paths of length *n*.

Call such a pattern recognition-and-response cognitive process ergodic. Not all cognitive processes are likely to be ergodic, implying that *H*, if it indeed exists at all, is path dependent, although extension to nearly ergodic processes, in a certain sense, seems possible (e.g., Wallace, 2005, pp. 31-32).

Invoking the Shannon-McMillan Theorem (Cover and Thomas, 2006; Khinchin, 1957), it becomes possible to define an adiabatically, piecewise stationary, ergodic information source **X** associated with stochastic variates *X*_*j*_ having joint and conditional probabilities *P*(*a*_0_*, …, a_n_*) and *P* (*a*_*n*_|*a*_0_, …, *a*_*n*−1_) such that appropriate joint and conditional Shannon uncertainties satisfy the classic relations

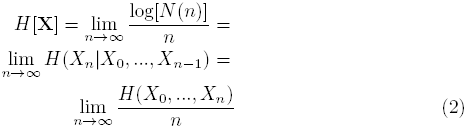

This information source is defined as dual to the underlying ergodic cognitive process.

‘Adiabatic’ means that, when the information source is properly parameterized, within continuous ‘pieces’, changes in parameter values take place slowly enough so that the information source remains as close to stationary and ergodic as needed to make the fundamental limit theorems work. ‘Stationary’ means that probabilities do not change in time, and ‘ergodic’ that cross-sectional means converge to long-time averages. Between pieces, as will be described below, it is necessary to invoke phase change formalism, a ‘biological’ renormalization that generalizes Wilson’s (1971) approach to physical phase transition (Wallace, 2005).

Shannon uncertainties *H*(*…*) are cross-sectional law-of-large-numbers sums of the form − *∑*_*k*_ *P*_*k*_ log[*P*_*k*_], where the *P*_*k*_ constitute a probability distribution. See Cover and Thomas (2006), Ash (1990), or Khinchin (1957) for the standard details.

We are not, however, constrained in this approach to the Atlan-Cohen model of cognition that, through the comparison with an internal picture of the world, invokes representation. The essential inference is that a broad class of cognitive phenomena – with and without representation – can be associated with a dual information source. The argument is direct, since cognition inevitably involves choice, choice reduces uncertainty, and this implies the existence of an information source.

Extension to non-ergodic information sources can be done using the methods of Wallace (2005, Sec. 3.1).

### 4 Groupoid symmetries

For cognitive systems, an equivalence class algebra can be constructed by choosing different origin points *a*_0_, and defining the equivalence of two states *a*_*m*_, *a*_*n*_ by the existence of high probability meaningful paths connecting them to the same origin point. Disjoint partition by equivalence class, analogous to orbit equivalence classes for a dynamical system, defines the vertices of a network of cognitive dual languages that interact to actually constitute the system of interest. Each vertex then represents a different information source dual to a cognitive process. This is not a representation of a network of interacting physical systems as such, in the sense of network systems biology (e.g., Arrell and Terzic, 2010). It is an abstract set of languages dual to the set of cognitive processes of interest, that may become linked into higher order structures.

Topology is now an object of algebraic study, so-called algebraic topology, via the fundamental underlying symmetries of geometric spaces. Rotations, mirror transformations, simple (‘affine’) displacements, and the like, uniquely characterize topological spaces, and the networks inherent to cognitive phenomena having dual information sources also have complex underlying symmetries: characterization via equivalence classes defines a groupoid, an extension of the idea of a symmetry group, as summarized by Brown (1987) and Weinstein (1996). Linkages across this set of languages occur via the groupoid generalization of Landau’s spontaneous symmetry breaking arguments that will be used below (Landau and Lifshitz, 2007; Pettini, 2007).

### 5 ‘Environment’ as an information source

Multifactorial cognitive and behavioral systems interact with, affect, and are affected by, embedding ‘environments’, in a large sense, that remember interaction by various mechanisms. It is possible to reexpress environmental dynamics in terms of a grammar and syntax that represent the output of an information source – another generalized language.

Obviously, within an organism, social assemblage of organisms, or ecosystem, different structures may, at different times, play the role of ‘system’ and of ‘environment’, of controllee and of controller.

Some examples:

1. The turn-of-the seasons in a temperate climate, for many ecosystems, looks remarkably the same year after year: the ice melts, the migrating birds return, the trees bud, the grass grows, plants and animals reproduce, high summer arrives, the foliage turns, the birds leave, frost, snow, the rivers freeze, and so on.
2. Human interactions take place within fairly well defined social, cultural, and historical constraints, depending on context: birthday party behaviors are not the same as cocktail party behaviors in a particular social set, but both will be characteristic.
3. Gene expression during development is highly patterned by embedding environmental context via ‘norms of reaction’ (e.g., Wallace and Wallace, 2010, 2013; Wallace, 2014).

Suppose it possible to coarse-grain the generalized ‘ecosystem’ at time *t*, in the sense of symbolic dynamics (e.g., Beck and Schlogl, 1993) according to some appropriate partition of the phase space in which each division *A*_*j*_ represent a particular range of numbers of each possible fundamental actor in the generalized ecosystem, along with associated larger system parameters. What is of particular interest is the set of longitudinal paths, system statements, in a sense, of the form *x*(*n*) = *A*_0_*, A*_1_*, …, A_n_* defined in terms of some natural time unit of the system. Thus *n* corresponds to an again appropriate characteristic time unit *T*, so that *t* = *T*, 2*T, …, nT*.

Again, the central interest is in serial correlations along paths.

Let *N*(*n*) be the number of possible paths of length *n* that are consistent with the underlying grammar and syntax of the appropriately coarsegrained embedding ecosystem, in a large sense. As above, the fundamental assumptions are that – for this chosen coarse-graining – *N*(*n*), the number of possible grammatical paths, is much smaller than the total number of paths possible, and that, in the limit of (relatively) large *n*, *H* = lim_*n*→∞_ log[*N*(*n*)]*/n* both exists and is independent of path.

These conditions represent a parallel with parametric statistics in that systems for which the assumptions are not true will require specialized approaches.

Nonetheless, not all possible ecosystem coarse-grainings are likely to work, and different such divisions, even when appropriate, might well lead to different descriptive quasi-languages for the ecosystem of interest. Thus, empirical identification of relevant coarse-grainings for which this theory will work may represent a difficult scientific problem.

Given an appropriately chosen coarse-graining, define joint and conditional probabilities for different ecosystem paths, having the form *P* (*A*_0_*, A*_1_*, …, A_n_*), *P* (*A*_*n*_|*A*_0_, …, *A*_*n*−1_), such that appropriate joint and conditional Shannon uncertainties can be defined on them that satisfy equation (2).

Taking the definitions of Shannon uncertainties as above, and arguing backwards from the latter two parts of equation (2), it is indeed possible to recover the first, and divide the set of all possible ecosystem temporal paths into two subsets, one very small, containing the grammatically correct, and hence highly probable paths, that we will call ‘meaningful’, and a much larger set of vanishingly low probability.

### 6 Interacting information sources

Given a set of cognitive modules (having dual information sources) that are linked to solve a problem, the ‘no free lunch’ theorem (English, 1996; Wolpert and Macready, 1995, 1997) extends a network theory-based theory (e.g., Arrell and Terzic, 2010). Wolpert and Macready show there exists no generally superior computational function optimizer. That is, there is no ‘free lunch’ in the sense that an optimizer pays for superior performance on some functions with inferior performance on others gains and losses balance precisely, and all optimizers have identical average performance. In sum, an optimizer has to pay for its superiority on one subset of functions with inferiority on the complementary subset.

This result is well-known using another description. Shannon (1959) recognized a powerful duality between the properties of an information source with a distortion measure and those of a channel. This duality is enhanced if we consider channels in which there is a cost associated with the different letters. Solving this problem corresponds to finding a source that is right for the channel and the desired cost. Evaluating the rate distortion function for a source corresponds to finding a channel that is just right for the source and allowed distortion level.

Yet another approach to the same result is the through the ‘tuning theorem’ (Wallace, 2005, Sec. 2.2), which inverts the Shannon Coding Theorem by noting that, formally, one can view the channel as ‘transmitted’ by the signal. Then a dual channel capacity can be defined in terms of the channel probability distribution that maximizes information transmission assuming a fixed message probability distribution.

From the no free lunch argument, Shannon’s insight, or the ‘tuning theorem’, it becomes clear that different challenges facing any cognitive system, distributed collection of them, or interacting set of other information sources, that constitute an organism must be met by different arrangements of cooperating modules represented as information sources.

It is possible to make a very abstract picture of this phenomenon based on the network of linkages between the information sources dual to the individual ‘unconscious’ cognitive modules (UCM), and those of related information sources with which they interact. That is, a shifting, task-mapped, network of information sources is continually reexpressed: given two distinct problems classes confronting the organism, there must be two different wirings of the information sources, including those dual to the available UCM, with the network graph edges measured by the amount of information crosstalk between sets of nodes representing the different sources.

Thus living systems involve interaction between very general sets of information sources assembled into a ‘task-specific device’ in the sense of Bingham (1988) that is necessarily highly tunable. This mechanism represents a broad generalization of the ‘shifting spotlight’ characterizing the global neuronal workspace model of consciousness (Baars, 1988; Wallace, 2005, 2012).

The mutual information measure of cross-talk is not inherently fixed, but can continuously vary in magnitude. This suggests a parameterized renormalization: the modular network structure linked by crosstalk has a topology depending on the degree of interaction of interest.

Define an interaction parameter *ω*, a real positive number, and look at geometric structures defined in terms of linkages set to zero if mutual information is less than, and ‘renormalized’ to unity if greater than, *ω*. Any given *ω* will define a regime of giant components of network elements linked by mutual information greater than or equal to it.

Now invert the argument: a given topology for the giant component will, in turn, define some critical value, *ω*_*C*_, so that network elements interacting by mutual information less than that value will be unable to participate, i.e., will be locked out and not be consciously or otherwise perceived. See Wallace (2005, 2012a) for details. Thus *ω* is a tunable, syntactically-dependent, detection limit that depends critically on the instantaneous topology of the giant component of linked information sources defining the analog to a global broadcast of consciousness. That topology is the basic tunable syntactic filter across the underlying modular structure, and variation in *ω* is only one aspect of more general topological properties that can be described in terms of index theorems, where far more general analytic constraints can become closely linked to the topological structure and dynamics of underlying networks, and, in fact, can stand in place of them (Atyah and Singer, 1963; Hazewinkel, 2002).

Given a cognitive system by an information source *X*, in the context of a set of ‘environmental’ information sources *Y*_1_*, …Y_k_*, we are particularly interested in the joint source defined by *H*(*X, Y*_1_*, …, Y_k_*), and next examine some details of how the mutually embedded system might operate in real time, focusing on the role of rapidly-changing feedback information via an extension of the Data Rate Theorem. That is, for an ‘organism’ (or identifiable subsystem of one) interacting with an ‘environment’ (in a large sense that may include other subsystems of that organism), we are interested in dynamics for which *ω* ∝ ***ℋ***.

### 7 Punctuated critical phenomena

The homology between the information source uncertainty dual to a cognitive process and the free energy density of a physical system arises from the formal similarity between their definitions in the asymptotic limit. Information source uncertainty can be defined as in the first part of equation (2). This is quite analogous to the free energy density of a physical system in terms of the thermodynamic limit of infinite volume (e.g., Wilson, 1971; Wallace, 2005). Feynman (2000) provides a series of physical examples, based on Bennett’s (1988) work, where this homology is an identity, at least for very simple systems. Bennett argues, in terms of idealized irreducibly elementary computing machines, that the information contained in a message can be viewed as the work saved by not needing to recompute what has been transmitted.

It is possible to model a cognitive system interacting with an embedding environment using an extension of the language-of-cognition approach above. Recall that cognitive processes can be formally associated with information sources, and how a formal equivalence class algebra can be constructed for a complicated cognitive system by choosing different origin points in a particular abstract ‘space’ and defining the equivalence of two states by the existence of a high probability meaningful path connecting each of them to some defined origin point within that space.

Recall that disjoint partition by equivalence class is analogous to orbit equivalence relations for dynamical systems, and defines the vertices of a network of cognitive dual languages available to the system: each vertex represents a different information source dual to a cognitive process. The structure creates a large groupoid, with each orbit corresponding to a transitive groupoid whose disjoint union is the full groupoid, and each subgroupoid associated with its own dual information source. Larger groupoids will, in general, have ‘richer’ dual information sources than smaller.

We can now begin to examine the relation between system cognition and the feedback of information from the rapidly-changing real-time environment, ***ℋ***, in the sense of equation (1).

With each subgroupoid *G*_*i*_ of the (large) cognitive groupoid we can associate a joint information source uncertainty *H*(*X*_*G*_*i*__, *Y* ) ≡ *H*_*G*_*i*__, where *X* is the dual information source of the cognitive phenomenon of interest, and *Y* that of the embedding environmental context.

Real time dynamic responses of a cognitive system can now be represented by high probability paths connecting ‘initial’ multivariate states to ‘final’ configurations, across a great variety of beginning and end points. This creates a similar variety of groupoid classifications and associated dual cognitive processes in which the equivalence of two states is defined by linkages to the same beginning and end states. Thus, we will show, it becomes possible to construct a ‘groupoid free energy’ driven by the quality of rapidly-changing, real-time information coming from the embedding ecosystem, represented by the information rate ***ℋ***, to be taken as a temperature analog.

The argument-by-abduction from physical theory is, then, that ***ℋ*** constitutes a kind of thermal bath for the processes of channeled cognition. Thus we can, in analogy with the standard approach from physics (Pettini, 2007; Landau and Lifshitz, 2007) construct a Morse Function by writing a pseudo-probability for the jointly-defined information sources *X*_*G*_*i*__, *Y* having source uncertainty *H*_*G*_*i*__ as

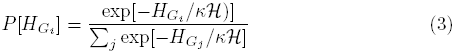

where *к* is an appropriate dimensionless constant characteristic of the particular system. The sum is over all possible subgroupiods of the largest available symmetry groupoid. Again, compound sources, formed by the (tunable, shifting) union of underlying transitive groupoids, being more complex, will have higher free-energy-density equivalents than those of the base transitive groupoids.

A possible Morse Function for invocation of Pettini’s topological hypothesis or Landau’s spontaneous symmetry breaking is then a ‘groupoid free energy’ *F* defined by

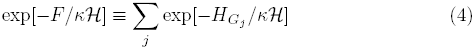

It is possible, using the free energy-analog *F*, to apply Landau’s spontaneous symmetry breaking arguments, and Pettini’s topological hypothesis, to the groupoid associated with the set of dual information sources.

Many other Morse Functions might be constructed here, for example based on representations of the cognitive groupoid(s). The resulting qualitative picture would not be significantly different.

Again, Landau’s and Pettini’s insights regarding phase transitions in physical systems were that certain critical phenomena take place in the context of a significant alteration in symmetry, with one phase being far more symmetric than the other (Landau and Lifshitz, 2007; Pettini, 2007). A symmetry is lost in the transition – spontaneous symmetry breaking. The greatest possible set of symmetries in a physical system is that of the Hamiltonian describing its energy states. Usually states accessible at lower temperatures will lack the symmetries available at higher temperatures, so that the lower temperature phase is less symmetric. The randomization of higher temperatures ensures that higher symmetry/energy states will then be accessible to the system. The shift between symmetries is highly punctuated in the temperature index.

The essential point is that decline in the richness of real-time environmental feedback ***ℋ***, or in the ability of that feedback to influence response, as indexed by *к*, can lead to punctuated decline in the complexity of cognitive process within the entity of interest, according to this model.

This permits a Landau-analog phase transition analysis in which the quality of incoming information from the embedding ecosystem – feedback – serves to raise or lower the possible richness of cognitive response to patterns of challenge. If *к****ℋ*** is relatively large – a rich and varied real-time environment – then there are many possible cognitive responses. If, however, noise or simple constraint limit the magnitude of *к****ℋ***, then behavior collapses in a highly punctuated manner to a kind of ground state in which only limited responses are possible, represented by a simplified cognitive groupoid structure.

These results represent a significant generalization of the Data Rate Theorem, as expressed in equation (1).

### 8 Renormalization

Certain details of information phase transitions can be calculated using ‘biological’ renormalization methods (Wallace, 2005, Section 4.2) analogous to, but much different from, those used in the determination of physical phase transition universality classes (Wilson, 1971).

Given *F* as a free energy analog, what are the transitions between adiabatic realms? Suppose, in classic manner, it is possible to define a characteristic ‘length’, say *r*, on the system, as described in the Mathematical Appendix. It is then possible to define renormalization symmetries in terms of the ‘clumping’ transformation, so that, for clumps of size *R*, in an external ‘field’ of strength *J* (that can be set to 0 in the limit), one can write, in the usual manner (e.g., Wilson, 1971)

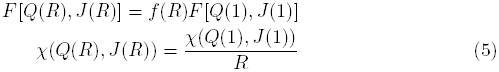

where *χ* is a characteristic correlation length and *Q* is an ‘inverse temperature measure’, i.e., ∝ 1/*к**ℋ***.

As described in Wallace (2005), very many ‘biological’ renormalizations, *f*(*R*), are possible that lead to a number of quite different universality classes for phase transition. Indeed, a ‘universality class tuning’ can be used as a tool for large-scale regulation of the system. While Wilson (1971) necessarily uses *f*(*R*) ∝ *R*^3^, following Wallace (2005), it is possible to argue that, since *F* is a kind of information measure, it is likely to ‘top out’ at different rates with increasing system size, so other forms of *f*(*R*) must be explored. Indeed, standard renormalization calculations for *f*(*R*) ∝ *R*^*δ*^, *m* log(*R*) + 1, and exp[*m*(*R −* 1)*/R*] all carry through.

### 9 Discussion and conclusions

The argument leading to equations (4) and (5) significantly extends the Data Rate Theorem of equation (1) in the context of cognitive systems, including those of a living state characterized – even defined – by cognition at every scale and level of organization. The unification of information theory and control theory via the Data Rate Theorem is not additive, but synergistic, encompassing a new body of statistical models that, in the sense of Dretske (1994), greatly constrain the possible dynamics of the living state. That is, in addition to providing important new conceptual approaches, this emerging body of theory may permit construction of new analytic tools that, like regression models, can provide benchmarks against which to compare experimental or observational data (e.g., Wallace, 2014b, Wallace and Wallace, 2014).

To put it another way, as has been long understood in robotics, walking across a crowded room, an exercise in embodied cognition, is far more difficult than playing a good game of chess (e.g., Brooks, 1991; Wilson and Golonka, 2013). Wallace (2012b) argues that a similar paradox exists at the cellular level, involving the many ‘glycosynapses’ (Cohen and Varki, 2010) operating at a literally astronomical number of cell surfaces in higher animals. The very concept of ‘information that controls the living state’ is, in fact, a misnomer, since the living state is cognitive at every scale and level of organization, and cognition, inherently involving active choices that reduce uncertainty, is characterized by nested, interacting information sources. This fact places necessary condition restrictions on the dynamics of life that are expressed in the Shannon Coding and Source Coding Theorems, the Rate Distortion Theorem, and the new Data Rate Theorem.

These are not trivial matters, either conceptually or formally. It took nearly half a century to unify information and control theories, and further development of useful mathematical tools is not likely to be less arduous. Even filling in the groupoid/Morse Theory outline presented here will not be easy. That outline, however, holds great promise across a spectrum of cognitive disciplines (e.g., Wallace, 2014c, d).

### 10 Mathematical appendix

In order to define the metric *r* used in equation (5), impose a topology on the system, so that, near a particular ‘language’ *A* defining some *H*_*G*_ there is (in an appropriate sense) an open set *U* of closely similar languages Â, such that *A*, *Â* ⊂ *U*.

Since the information sources are ‘similar’, for all pairs of languages *A, Â* in *U*, it is possible to:

1. Create an embedding alphabet which includes all symbols allowed to both of them.
2. Define an information-theoretic distortion measure in that extended, joint alphabet between any high probability (grammatical and syntactical) paths in *A* and *Â*, written as *d*(*Ax*, *Âx*) (Cover and Thomas, 2006). Note that these languages do not interact in this approximation.
3. Define a metric on *U*, for example,

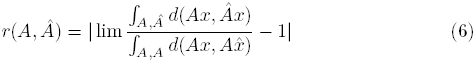

using an appropriate integration limit argument over the high probability paths. Note that the integration in the denominator is over different paths within *A* itself, while in the numerator it is between different paths in *A* and *Â*. Consideration suggests *r* is indeed a formal metric.

Other approaches to metric construction on *U* are possible, as are other approaches to renormalization and phase transition.

